# Ventral hippocampus mediates the context-dependence of two-way signaled avoidance in male rats

**DOI:** 10.1101/2021.02.04.429792

**Authors:** Cecily R. Oleksiak, Karthik R. Ramanathan, Olivia W. Miles, Sarah J. Perry, Stephen Maren, Justin M. Moscarello

## Abstract

Considerable work indicates that instrumental responding is context-dependent, but the neural mechanisms underlying this phenomenon are poorly understood. Given the important role for the hippocampal formation in contextual processing, we hypothesized that reversible inactivation of the hippocampus would impair the context-dependence of active avoidance. To test this hypothesis, we used a two-way signaled active avoidance (SAA) task that requires rats to shuttle across a divided chamber during a tone CS in order to avoid a footshock US. After training, avoidance responding was assessed in an extinction test in both the training context and a novel context in a counterbalanced order. Rats performed significantly more avoidance responses in the training context than in the novel context, demonstrating the context-dependence of shuttle avoidance behavior. To examine the role of the hippocampus in the context-dependence of SAA, we reversibly inactivated either the dorsal (DH) or ventral hippocampus (VH) prior to testing. Inactivation of the VH eliminated the context-dependence of SAA and elevated avoidance responding in the novel context to levels similar to that expressed in the training context. In contrast, DH inactivation had no effect on avoidance in either context, and neither manipulation affected freezing behavior. Therefore, the integrity of the VH, but not DH, is required for the expression context-dependence of avoidance behavior.

## 1 INTRODUCTION

Coping with complex threats in a fluid environment requires the ability to bypass innate behavioral reactions and take action in order to avoid harm (Moscarello & Hartley, 2017; Moscarello & Maren, 2018). Such proactive defensive strategies are beneficial to the extent that they protect against real-world danger. However, even an adaptive avoidance behavior can have a highly disruptive effect on normal life activities when it is triggered in a safe situation that does not involve the relevant threat (Craske et al., 2017; Lovibond, Mitchell, Minard, Brady, & Menzies, 2009; Pittig, Wong, Glück, & Boschet, 2020). Thus, to prevent the excessive, maladaptive expression of defensive action, proactive avoidance strategies must be confined to the contexts in which the danger they defend against is imminent and real.

In order to explore the relationship between context and the expression of avoidant behavior, we employed a two-way signaled active avoidance (SAA) paradigm for rats in which the subject learns to shuttle across a divided chamber during a conditioned stimulus (CS, tone) in order to avoid an aversive unconditioned stimulus (US, shock). SAA training is a multiphase learning process that involves the rapid formation of a Pavlovian memory followed by the acquisition of an avoidance contingency. When a sequential form of learning produces multiple memories related to the same cue, the first memory acquired tends to be elemental and can be reactivated in multiple environments, whereas the second memory often requires the disambiguating influence of context in order to be successfully retrieved (Bouton, 1993; Bouton, 2004). Thus, contextual stimuli may function to constrain the expression of the avoidance response to the environment in which SAA training occurs, creating a psychological mechanism that prevents an adaptive form of avoidance from generalizing to situations in which it may prove disruptive.

The hippocampus plays an important role in the contextual modulation of memory retrieval (Fanselow & Dong, 2010; Holland & Bouton, 1999; Maren, Phan, & Liberzon, 2013). Though the dorsal (DH) and ventral (VH) subdivisions of the hippocampus are thought to have distinct functions (Fanselow & Dong, 2010), both DH and VH have been demonstrated to mediate the renewal of CS-evoked freezing after extinction (Ji & Maren, 2005; Marek et al., 2018; Maren & Hobin, 2007; Xu et al., 2016), suggesting that both regions participate in the process by which contextual information regulates the expression of cued aversive memory. Consistent with this, work in rabbits has shown that the context-dependent expression of SAA in a running wheel apparatus is disrupted by lesions of the entorhinal cortex (Freeman, Weible, Rossi, & Gabriel, 1997), a key cortical interface between hippocampus and other brain regions (Fanselow & Dong, 2010). On this basis, we hypothesized that the hippocampus limits instrumental avoidance to the training context by suppressing this response outside the SAA training context, thus functioning to apply an adaptive constraint to the expression of avoidant behavior.

The experiments described here demonstrate that, after the acquisition of the avoidance response (shuttling), exposure to the CS in a novel shuttle box context causes a decrease in avoidance and an increase in CS-evoked freezing, suggesting a retrieval of the first-learned Pavlovian memory in the absence of the contextual cues needed for the expression of avoidance. We then demonstrate that muscimol inactivation of VH, but not DH, prevents suppression of the avoidance response in a novel context, thereby eliminating the context-dependent expression of this behavior. These data confirm that hippocampal circuits constrain the expression of avoidance in novel environments and demonstrate that the contextually appropriate expression of an avoidance strategy is regulated by a VH-dependent mechanism.

## 2 METHODS

### 2.1: Subjects

Subjects were 111 experimentally naïve, adult male Sprague Dawley rats (200-300 g) purchased from Envigo. From their arrival, the rats were kept in a vivarium that was temperature/humidity controlled on a 14h light: 10h dark cycle beginning at 7 AM with free access to food and water in individual cages. Experiments were conducted during the light part of the cycle. Experimenters handled the rats for 2 minutes/day for around 5 days to habituate animals to being held before surgeries or behavior. All experiments were approved by the Animal Care and Use Committee from Texas A&M University.

### 2.2: Surgical procedure

One week prior to training, rats were anesthetized with isoflurane (5% for induction, slowly reduced throughout procedure) and their heads secured and leveled in a stereotaxic apparatus (Kopf Instruments). An incision was made down the center of the scalp, bregma was identified, and a drill was used to make holes for 3-4 jewelers screws. Holes were also made for bilateral guide cannulae (either 11 or 5 mm, 26 gauge, Plastics One) targeting either the ventral hippocampus (A/P: −5.25, M/L: ± 5, D/V: −7.3 from bregma) or dorsal hippocampus (A/P: −3.5, M/L: ± 2.4, D/V: −3.5 from bregma). The guide cannulae were secured to the skull using dental cement. Cannula dummies made of stainless steel (30 gauge, 12 or 6 mm, Plastics One) were used to protect guide cannula from blockage. Rats were given 7 days to recover in which the dummies were changed twice to habituate animals to infusion procedures.

### 2.3: Behavioral apparatus

Four shuttle boxes (50.8 × 25.4 × 30.5 cm, LXWXH; Coulbourn Instruments) constructed of Plexiglas and metal were used for all two-way signaled active avoidance tasks. The shuttle boxes were divided into two compartments by a metal divider with an open passage (8 × 9 cm, WXH) to allow animals to cross freely between the compartments. The floors consisted of conductive stainless-steel bars through which electric shock was delivered. Two speakers, one integrated into each wall opposite the divider, delivered a 2 kHz, 80 dB tone CS for 15 seconds. A scrambled shocker delivered a 0.7 mA, 0.5 second footshock US through the floor. Each shuttle box was contained inside a larger sound-attenuating chamber. The subjects’ motion was monitored by two infrared arrays, one in each compartment in order to detect shuttling (crossing from one compartment to the other in either direction).

In order to examine the role of context in SAA, we trained rats in two distinct shuttle box environments, Context A and Context B. Context A had an illuminated house light in each compartment (0.5 W light bulb), black-and-white striped wall inserts, a 3% acetic acid odor, and the doors of the sound-attenuating chamber remained closed during training. Rats were transported to Context A in white transport boxes with bedding. Context B had no house light or compartment lights, black construction paper wall inserts with glow-in-the-dark stars, a 1% ammonia odor, and the doors of the sound-attenuating chamber remained open during training (room lights also remained off). Rats were transported to Context B in black transport boxes without bedding. Whenever the rats were tested in the context that differed from the one in which they were trained (i.e., context B for the groups trained in A and Context A for the groups trained in B), a black Plexiglas floor insert was added to create a texture cue that distinguished the test context from the one in which training occurred.

### 2.4: Drug infusions

For VH and DH infusions, rats were carried to a separate room in the laboratory using white 5-gallon buckets with bottoms covered in bedding. After dummy removal, stainless steel injectors (33 gauge, 12 or 6 mm respectively) were fit into the guide cannulae. These injectors were connected to polyethylene tubing, which attached to 10-µl Hamilton syringes. The syringes were affixed to an infusion pump (Kd Scientific). Tubing was filled with distilled water, an air bubble was pulled up, and then drug or vehicle was filled to the desired amount. Muscimol (diluted to 0.1 µg/µl with sterile saline) was infused at a rate of 0.1 µl/min for 5 minutes for the VH (for a total dose of .05 µg/0.5 µl) and 3 minutes for DH (for a total dose of 0.03 µg/0.3 µl). Injectors were left to sit for 2-3 minutes and air bubble movement was monitored to ensure infusion. Then new dummies were used to cap the guides, and animals were conveyed to the behavior room in the contextually appropriate transport box.

### 2.5: Signaled active avoidance training

During the first session of SAA training, rats were presented with a single Pavlovian trial in which the CS was paired with the US in a way that could not be avoided, in order to ensure that the tone would be associated with the shock. The rest of training consisted of avoidance trials in which an avoidance response (shuttling during the CS) would result in the immediate offset of the CS and the omission of the US. The rats received 30 avoidance trials during each SAA training session. Between each trial was an ISI that averaged 120 seconds. CS and US presentation were controlled by GraphicState Software (Coulbourn Instruments), which also collected shuttling data via the infrared arrays in each compartment. Rats were considered poor avoiders and excluded from the statistical analyses if they avoided on ≤ 20% of trials for the last 3 days of training.

### 2.6: Signaled active avoidance testing under extinction conditions

Animals received counterbalanced tests in both the Training Context and in a Novel Context in which they were not trained (e.g., if a subject was trained in A then B was novel and vice versa). During these tests, ten 15-sec CSs were presented under extinction conditions, meaning that the CS did not terminate with shuttling and no US was presented. This design created an unbiased, within-subjects test of the effects of context on SAA performance.

### 2.7: Experiments

#### 2.7.1: The role of context in two-way signaled active avoidance

Our first experiment assessed the role of context in SAA at two different training time points. To do this, rats were given 4 days of SAA training (the normal time to asymptote in our paradigm; Ramirez, Moscarello, LeDoux, & Sears, 2015) in one of two training environments, Context A or Context B. This was followed by 2 days of testing under extinction conditions in which rats were exposed to the CS in the Training and Novel contexts in a counterbalanced order. We followed this with 4 more days of SAA in the Training Context and then 2 additional days of counterbalanced tests in the Training and Novel contexts.

#### 2.7.2: The role of ventral and dorsal hippocampus in the context-dependence of signaled active avoidance

Our second and third experiments examined whether the effect of context on SAA is mediated by VH or DH. Because our first experiment revealed a comparable context shift effect after both 4 and 8 days of SAA training, we limited training to 4 days in both of these experiments. In addition, because the strongest effects in our first experiment occurred when we used Context B for Training and Context A for Novel, we chose to focus on this training and test arrangement for our manipulations of VH and DH. Thus, in both experiments, animals were trained for 4 days in Context B prior to tests in both Context B (Training) and Context A (Novel) delivered in a counterbalanced order. Rats were given either muscimol or vehicle infusions into either VH or DH (depending on the experiment) approximately 10 minutes before both training and novel tests. Assignment to muscimol or vehicle conditions, which occurred between the end of avoidance training and test, was done to ensure groups that were matched in terms of their average daily avoidance responses across the 4 days of SAA training.

### 2.8: Histology

Rats in drug experiments were injected with a fatal dose of sodium pentobarbital (Fatal Plus; 100 mg/ml, 0.7 ml) and were transcardially perfused first with refrigerated saline and then with 10% formalin. The brains were collected and were left in 10% formalin for 24 hours. The brains were then transferred to 30% sucrose in PBS at 4°C until they sank. Brains were sliced coronally on a cryostat (−20°C) and mounted on subbed slides and stained with thionin (0.25% thionin) to visualize cannula tracts.

### 2.9: Data analysis

In all three experiments, the acquisition of avoidance was quantified using the total number of responses made per SAA training session. For all tests conducted under extinction conditions, the number of avoidance responses was capped at 1 per CS for a total of 10 possible per test. To directly compare the impact of VH and DH inactivation on the effect of context shift, difference scores were calculated for avoidance responses during tests in the Training and Novel Contexts as follows: Novel/(Training+Novel). In addition, each test was recorded with a digital camera, and freezing during the CS was scored by a trained rater blinded to group. These data are presented as the percent time spent freezing averaged across the 10 CSs.

ANOVAs followed by the appropriate *post hoc* tests were performed on these data. All statistical analyses were run in Statview version 5.0.1 (SAS Institute) in a MacOS 9 open-source emulator.

## 3 RESULTS

### 3.1: Signaled active avoidance is context-dependent

Of the 32 rats that began the experiment, one rat was excluded due to poor avoidance and three were included in the avoidance data but excluded for freezing data due to video loss. To assess the effect of context on avoidance, we trained the rats for eight days in a two-way signaled active avoidance task, with a pair of tests consisting of 10 tones presented under extinction conditions after four days and another after eight days (Fig. 1). Avoidance responses over the course of the eight days of SAA training were analyzed using a two-way repeated measures ANOVA with the within-subjects factor of Day and the between-subjects factor of Training Environment (trained in either A or B). The ANOVA demonstrated a main effect of Day [F(7,203) = 41.95, p < 0.0001] and an interaction between Day and Training Environment [F(7,203) = 2.60, p = 0.014]. Thus, subjects learned the avoidance response in both A and B, but those trained in A performed significantly fewer responses over training (Fig. 2A).

**Figure 1.**
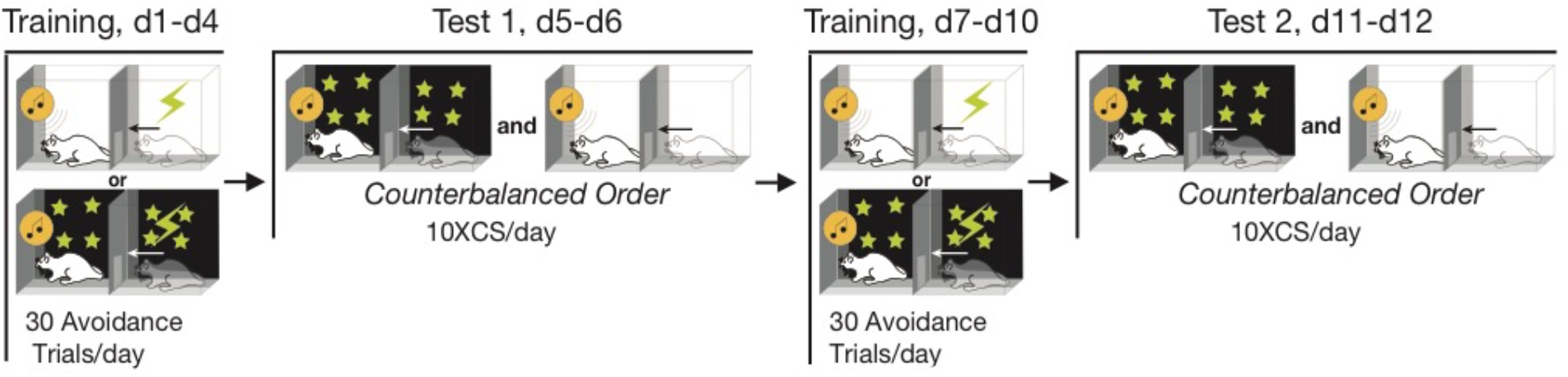
Behavioral design. Rats received 4 days of two-way signaled active avoidance (SAA) training in one of two distinct contexts, A or B. Subjects then received two daily tests conducted in either the Training or Novel contexts in a counterbalanced order. In each test, 10 CSs were presented under extinction conditions (no shock was delivered, and the CS did not inactivate if the subject performed an avoidance response). This was followed by 4 more days of SAA in each subject’s original training context and two final tests under extinction conditions in Training and Novel contexts.

**Figure 2.**
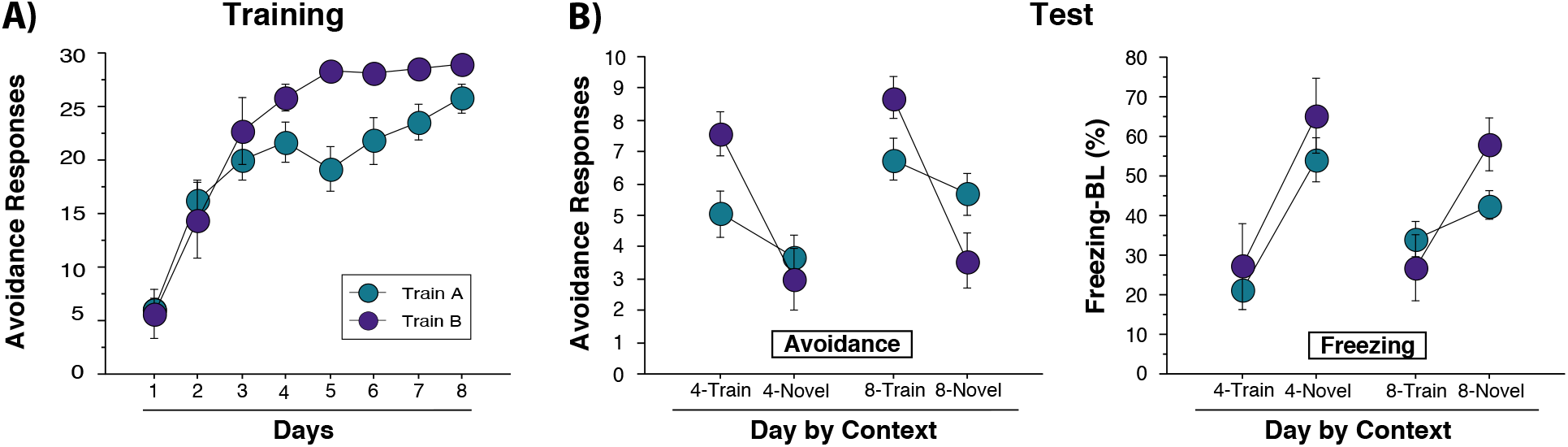
SAA is context-dependent after both 4 and 8 days of training. (A) Average number of avoidance responses performed on each day of SAA training. Rats trained in context A (n=24) performed significantly fewer avoidance responses than those trained in B (n=7). (B) (Left) Avoidance responses during counterbalanced tests conducted in Training and Novel contexts – in each, 10 CSs were delivered under extinction conditions and avoidance responses were capped at 1/CS for a possible total of ten; (Right) CS-evoked freezing during counterbalanced tests in Training and Novel contexts, expressed as a percentage of the 15-sec tone with baseline freezing levels subtracted out. Rats performed significantly fewer avoidance responses in the Novel context, though this effect was less pronounced in subjects trained in Context A. There was also a significant increase in freezing in the Novel context. All data are shown as mean ± SEM.

Tests were performed in both contexts in a counterbalanced manner. Capped avoidances (one response/CS) were assessed with a three factor ANOVA of mixed design with two within-subject factors of Test Context (Training or Novel) and Day (four or eight days), and a between-subjects factor of Training Environment (A or B). This revealed a main effect of Test Context, which indicated that all subjects performed significantly better in the Training context than in the Novel context [F(1,29) = 44.49, p < 0.0001]. However, we also found a significant Test Context by Training Environment interaction [F(1,29) = 15.81, p = 0.0004], indicating the effect of Test Context on avoidance responses was smaller in subjects trained in A (Fig 2B, Left). The ANOVA also revealed a main effect of Day [F(1,29) = 7.02, p = 0.013], but no significant Test Context by Day interaction [F(1,29) = 0.032, p = 0.86]. Thus, the effect of context on the avoidance response was comparable after both four and eight days of training.

Freezing evoked by the CS during all tests were analyzed with a three-factor ANOVA identical to that performed on avoidance data. This revealed a main effect of Test Context on freezing [F(1,26) = 34.09, p < 0.0001], but no interaction between Test Context and Training Environment [F(1,26) = 2.26, p = 0.15]. Thus, freezing was significantly higher in the Novel context compared to the Training context (Fig. 2B, Right). Therefore, the overall effect of testing outside the training context was a decrease in avoidance accompanied by an increase in freezing.

### 3.2: Ventral hippocampus inactivation reverses context-dependence of signaled active avoidance

#### 3.2.1: Histological analysis

To examine the role of VH in the context-dependence of avoidance, rats were implanted with cannula for future muscimol injections. Cannula placements for all animals included are presented in Figure 3A. Of the 43 animals that started the experiment, seven animals were excluded due to dislodged implants, seven were excluded to inaccurate placements, and four were excluded for poor avoidance. This led to a final grouping of 13 muscimol animals and 12 vehicle animals.

**Figure 3.**
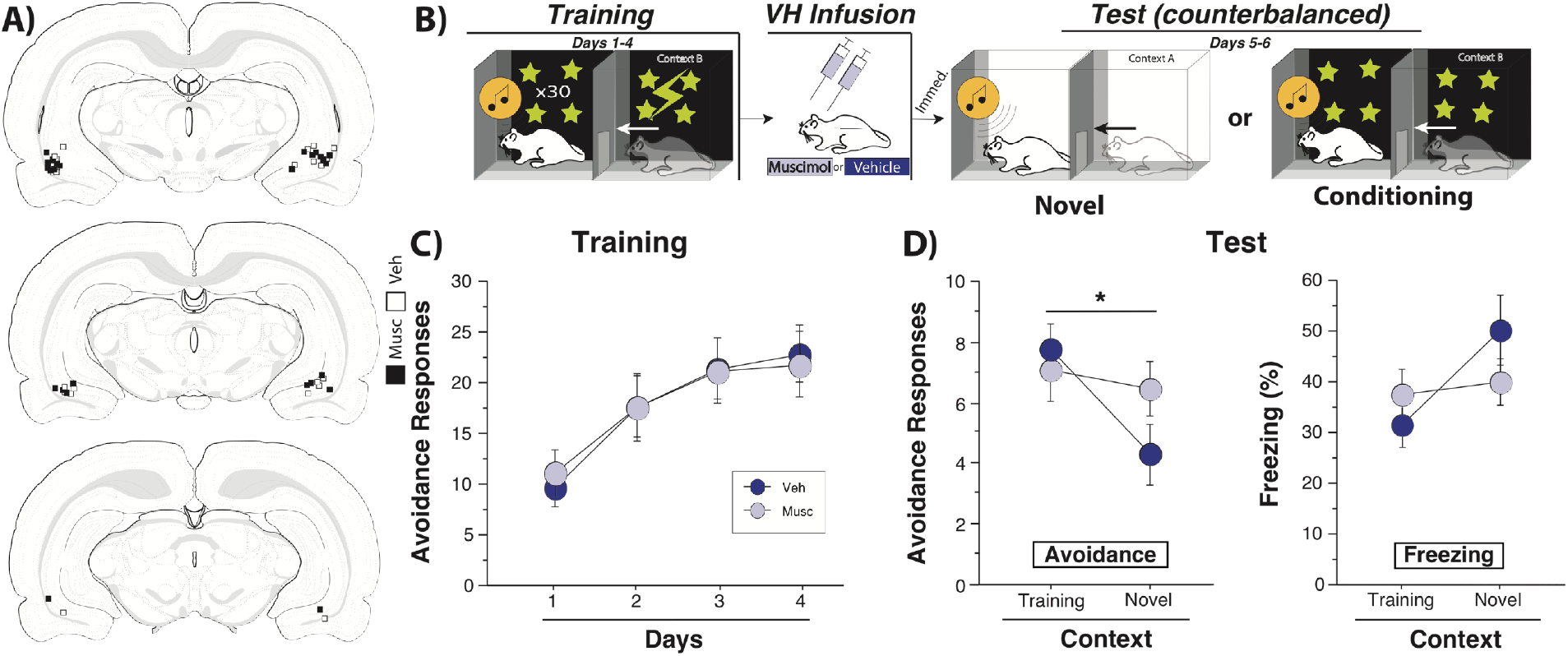
Ventral hippocampus (VH) mediates the effect of context on SAA. (A) Atlas images with cannula placements for all subjects included in the analysis (images from: Swanson, 1998) (B) Rats received 4 days of SAA training in context B prior to counterbalanced tests conducted under extinction conditions (as described in Fig. 1) in Training (B) and Novel (A) contexts. (C) Average number of avoidance responses per day for muscimol and vehicle groups. (D) Rats received either muscimol (n=13) or vehicle (n=12) infused into the VH immediately before each test; (Left) The effect of muscimol and vehicle on avoidance responses (capped at one/CS) during tests in Training and Novel contexts; (Right) the effect of muscimol and vehicle on CS-evoked freezing. VH inactivation caused rats to perform significantly more avoidance responses in the Novel) context compared to vehicle controls (* indicates p<0.05). All data presented as means ± SEM.

#### 3.2.2: Behavioral data

After recovery from surgery, subjects implanted with VH cannula received four days of SAA training in Context B before counterbalanced tests in A and B. Muscimol or vehicle was infused into VH prior to each test (Fig. 3B). Avoidance responses from training were analyzed with a mixed-design, two-way ANOVA with a within-subjects factor of Day (1-4) and a between-subjects factor of Drug Assignment (muscimol or vehicle). This analysis revealed a main effect of Day [F(3,69) = 49.403, p = 0.0001], but no effect for Drug Assignment [F(3,69) = 0.859, p = 0.47], suggesting that both groups acquired the avoidance response to the same level (Fig. 3C).

Avoidance responses from the test sessions were analyzed by a two-way ANOVA of mixed design with a within-subjects factor of Test Context (training or novel) and between-subjects factor of Drug (muscimol or vehicle). This analysis revealed an interaction between Test Context and Drug [F(1,23) = 7.150, p = 0.0136), indicating that the vehicle group demonstrated context-dependent avoidance, but the muscimol group performed similar number of responses in both the training and novel contexts (Fig 3D, Left). Planned comparisons confirmed the interaction, demonstrating that vehicle animals showed significantly reduced responding in the novel context relative to the training context [t(11) = 4.535, p = 0.0009]. In contrast, muscimol animals produced similar levels of avoidance in novel and training contexts [t(12) = 0.410, p = 0.6888]. Thus, temporary inactivation of VH made the avoidance response context-independent.

Freezing data from these tests were analyzed with a two-way, mixed-design ANOVA with a within-subjects factor of Test Context (training or novel) and between-subjects factor of Drug (muscimol or vehicle). This analysis revealed a main effect of Test Context [F(1,23) = 5.530, p = 0.0276], but no Test Context by Drug interaction [F (1,23) = 3.087, p = 0.0922], suggesting that VH inactivation does not impact freezing (Fig. 3D, Right).

To assess the effect of VH inactivation on baseline locomotion, we analyzed shuttling during the acclimation and inter-stimulus interval (ISI) period of test. We observed no drug assignment differences for VH for acclimation [F(1,23) = 0.0004, p = 0.985] or ISI [F(1,23) = 0.096, p = 0.759]. These data suggest that differences in locomotion due to the drug were not responsible for the increase in avoidance. Hence, these data demonstrate that muscimol ‘rescues’ avoidance responding in the novel context.

### 3.3: Dorsal hippocampus is not responsible for the context-dependence of signaled active avoidance

#### 3.3.1: Histological analysis

We next examined the role of DH in the context-dependence of SAA, rats were implanted with cannula in the DH. Cannula placements for all animals included are presented in Figure 4A. Of the 36 animals that started the experiment, two were excluded due to inaccurate placements, two were excluded due to incorrect infusions, and six were excluded due to poor avoidance. This led to a final grouping of 14 muscimol animals and 12 vehicle animals.

**Figure 4.**
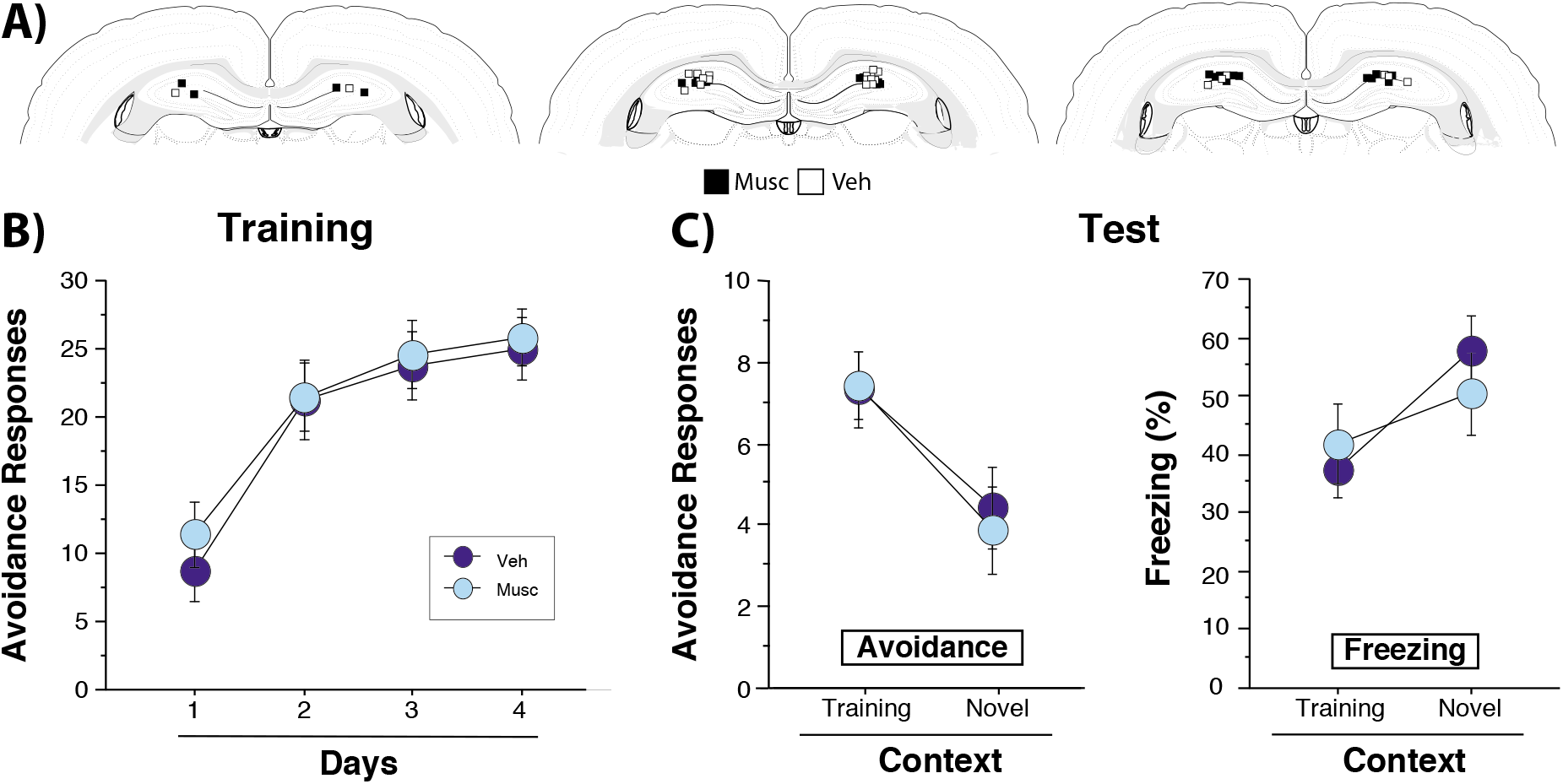
Dorsal hippocampus (DH) has no role in the context-dependence of SAA. (A) Atlas images with cannula placements for all muscimol (n=14) and vehicle (n=12) subjects included in the analysis (images from: Swanson, 1998). (B) Rats received 4 days of SAA training in context B prior to counterbalanced tests conducted under extinction conditions (as described in FIG 1) in Training (B) and Novel (A) contexts. (C) Rats received either muscimol (n=13) or vehicle (n=12) infused into the DH immediately before each test; (Left) The effect of muscimol and vehicle on avoidance responses (capped at 1/CS) during tests in Training and Novel contexts; (Right) the effect of muscimol and vehicle on CS-evoked freezing. DH muscimol animals show a similar reduction in avoidance responding and increase in freezing as vehicle controls. All data calculated as means ± SEM. DH inactivation had no effect on avoidance or freezing in either Training or Novel contexts.

#### 3.3.2: Behavioral analysis

After recovery from surgery, subjects implanted with DH cannula received four days of SAA training in Context B before counterbalanced tests in A and B. Avoidance responses from training were analyzed with a mixed-design, two-way ANOVA with a within-subjects factor of Day (1-4) and a between-subjects factor of Drug Assignment (muscimol or vehicle). This analysis revealed a main effect of Day [F (3,72) = 45.956, p = 0.0001], but no effect for Drug Assignment [F (3,72) = 0.244, p = 0.87], suggesting that both groups acquired the avoidance response to the same degree (Fig. 4B).

Muscimol or vehicle were infused into DH prior to each test. Avoidance responses from the test sessions were analyzed by a two-way ANOVA of mixed with a within-subjects factor of Test Context (training or novel) and between-subjects factor of Drug (muscimol or vehicle). There was a main effect of Test Context [F(1,24) =22.80, p < 0.0001] but no interaction between Test Context and Drug [F(1,24) = 0.232, p = 0.63), indicating that both groups performed fewer avoidance responses in the novel test context (Fig. 3C, Left). There was also a main effect of Test Context for freezing percentages [F(1,24) = 21.11, p = 0.0001] but no interaction between Test Context and Drug [F (1,24) = 3.378, p = 0.0785] similar to the VH animals (Fig. 4C, Right). Therefore, dorsal hippocampus is not involved in the context-dependence of SAA.

### 3.4: Comparison of ventral and dorsal hippocampus inactivation

To directly compare the results of the VH and DH inactivation experiments, we computed a difference score for avoidance responses in Training and Novel contexts for animals in the VH muscimol group, DH muscimol group, and a combined VH and DH vehicle group from both experiments (Fig. 5). A factorial ANOVA revealed a significant main effect of group [F(2,47) = 5.464, p = 0.0073]. A Fisher’s PLSD post-hoc test revealed that the VH muscimol group had significantly higher difference scores than both the DH muscimol group (p = 0.0031) and the vehicle animals (p= 0.0100). Thus, the effect of a context shift on avoidance is not observed under conditions of VH inactivation, whereas DH inactivation and vehicle treatments have no effect on the context dependence of avoidance.

**Fig. 5.**
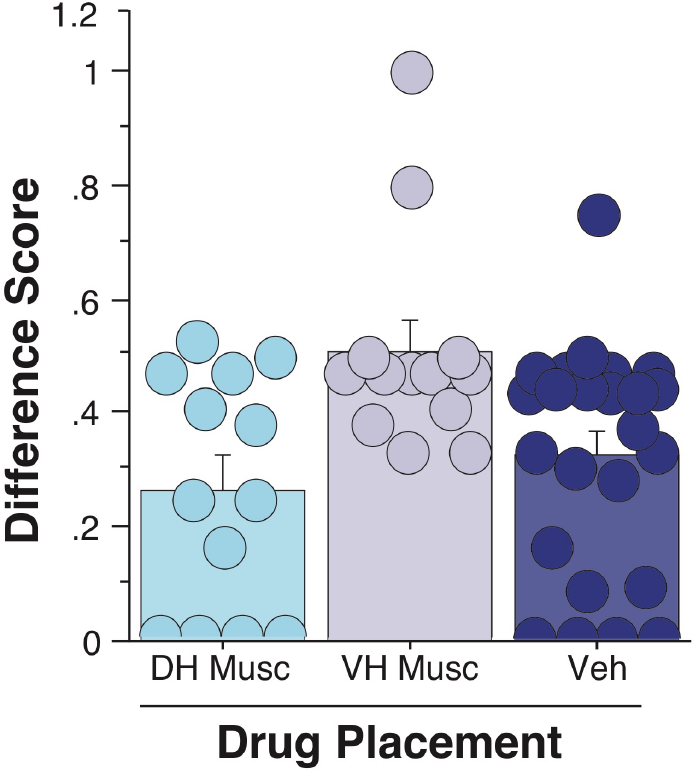
Difference scores calculated from avoidance responses during tests in Training and Novel contexts for DH muscimol, VH muscimol, and combined DH and VH vehicle groups. A score of 0-0.5 suggests more avoidance responses in the Training context, 0.5 indicates equal levels of avoidance in each, and 0.5- 1 signifies more responses in the Novel context. Subjects in the VH muscimol group performed significantly more responses in the Novel context relative to both the DH muscimol and vehicle groups. Data is represented as means + SEM.

## 4 DISCUSSION

Here, we present data on the role of context in two-way signaled active avoidance (SAA) behavior. We found that rats froze more and avoided less when the CS was presented in a novel context that differed from the experimental environment in which SAA training occurred, demonstrating the context-dependence of instrumental avoidance. Muscimol inactivation of the ventral hippocampus (VH) abolished the context-dependence of avoidance, restoring avoidance responding in a novel context to levels comparable to those observed in the training context. VH inactivation also caused a nonsignificant decrease in freezing in the novel context. Though VH muscimol was sufficient to eliminate the context-dependence of avoidance, inactivation of the dorsal hippocampus (DH) had no effect on avoidance or freezing. Thus, we conclude that two-way SAA behavior is context-dependent, and that VH functions to attenuate the avoidance response outside the context in which it is acquired.

Our behavioral results are highly consistent with prior work by Gabriel and colleagues demonstrating that rabbits trained in a wheel-running SAA task perform fewer responses in a novel context (Freeman et al., 1997). This group has argued that the effect of context on avoidance is mediated by a circuit consisting of entorhinal cortex⟶ventral subiculum⟶nucleus accumbens projections, which they termed the “ESA pathway” (Burhans & Gabriel, 2007). These regions are all highly connected (Groenewegen, Vermeulen-Van der Zee, te Kortschot, & Witter, 1987; Groenewegen, Wright, Beijer, & Voorn, 1999; van Groen, van Haren, Witter, & Groenewegen, 1986; Witter & Groenewegen, 1990), and, given that our VH manipulations included ventral subiculum, a similar pathway may indeed play a role in the context-dependence of two-way SAA. Indeed, rabbits with ventral subiculum lesions produce more wheel-turn avoidance responses in a novel context (Burhans & Gabriel, 2007). However, a notable difference between that report and the current study is that Buhrans & Gabriel (2007) demonstrate that ventral subiculum lesions reduced the expression of avoidance responses in the original training environment, whereas we show no effect of VH inactivation in the training context. This difference may be attributable to the contrasting effects of permanent lesion versus temporary inactivation, or to important differences in the task itself (i.e. wheel-running vs two-way SAA), or a combination of both.

Our initial experiment makes it clear that the sensory features of the different contexts had an impact on both the acquisition of avoidance and the magnitude of the decrement in avoidance responding outside the training context. Training in Context A produced a somewhat lower avoidance asymptote relative to Context B, and training in Context B followed by a shift to Context A produced a greater reduction in avoidance responding than a shift from Context B to A. One key difference between the two environments was that Context A was illuminated whereas Context B was dark. The nonassociative, attenuating effect of light on the locomotor behavior of rats (Crawley, 1985; File, 1980) could account for the relatively lower levels of the avoidance response observed in Context A. Because context shift effects were observed in both A and B, we conclude that nonassociative features of the sensory environment modulated but did not fully determine the effect of context on avoidance. Additionally, these effects were observed at two training time points, suggesting that novelty alone does not account for the context-shift deficit.

Memory interference occurs when two distinct responses to a single cue are acquired across sequential phases of a conditioning experiment, causing one response to occlude the other (Bouton, 1993). From this perspective, the avoidance response, which is acquired second, can be said to retroactively interfere with freezing, which is acquired first. Expression of the second response learned requires presentation of the relevant cue in conjunction with disambiguating background information, making retroactive interference context-dependent (Bouton, 2004). This is consistent with suppression of avoidance in a novel shuttle box context, which is associated with a return to CS-elicited freezing behavior. Because avoidant behavior becomes disruptive when it is triggered in neutral or safe situations (Craske et al., 2017; Lovibond et al., 2009; Pittig et al., 2020), the contextual requirements of retroactive interference may function as a psychological mechanism that constrains avoidance to scenarios in which it serves as an adaptive defense against danger. Thus, the balance between healthy and pathological avoidance strategies may indeed be determined by the degree to which interference phenomena limit avoidant behavior to specific situations and places.

VH inactivation caused an increase in the expression of the avoidance response in a novel context without producing a significant decrease in conditioned freezing, suggesting that the VH may be an important brain mechanism that limits the expression of the avoidance response to appropriate contexts. Cued extinction paradigms also involve the acquisition of sequential, opposing associations with the same CS, and thus are subject to retroactive interference and context-dependent memory expression (Bouton, 1993; Bouton, 2004; Bouton, Maren, & McNally, 2020). Indeed, the VH has been demonstrated to inhibit the retrieval of extinction memories when an extinguished CS is encountered in a different environment than the one in which extinction training occurred (Hobin, Ji, & Maren, 2006; Marek et al., 2018). Thus, in the case of both two-way SAA and cued extinction learning, the VH seems to enforce the contextual requirements of retroactive interference by attenuating the expression of the second memory acquired when the relevant cue is encountered outside the appropriate environment.

This characterization is broadly compatible with the idea that the VH is a key component of a ‘behavioral inhibition system’ (BIS). In the BIS framework, VH is recruited by situations that produce conflict between mutually exclusive behavioral responses, which VH resolves by facilitating ‘passive’ reactions such as freezing (Gray, 1982; Gray & McNaughton, 2000). Though the original formulation of the BIS describes the hippocampal-septal system as a substrate of anxiety created by behavioral conflict (Fernández-Teruel & Tobeña, 2020; Gray & McNaughton, 2000), a modern update of the theory, which incorporates the role of hippocampus in contextual processing (Bryant & Barker, 2020), is consistent with the evidence presented here indicating that VH regulates response conflict by suppressing contextually inappropriate memories and responses.

An alternative interpretation of our results is that VH inactivation impaired subjects’ ability to discriminate between the training and test contexts, resulting in similar and high levels of avoidance performance in both contexts. Though DH underpins context discrimination (Frankland, Cestari, Filipkowski, McDonald, & Silva, 1998; Wang, Teixeira, Wheeler, & Frankland, 2009), prior work has shown that VH inactivation does not have an impact on contextual discrimination (Hobin et al., 2006). The fact that we observed no effect for DH inactivation is intriguing, given that both contexts were necessarily similar enough in size, shape, and layout to allow shuttling to occur. This may be attributable to the repeated exposure animals received to the training context as a part of SAA training, consistent with evidence that repeated conditioning experiences in a single environment produce a more robust form of contextual discrimination (Wang et al., 2009).

In summary, the data presented here demonstrate the contextual dependence of avoidance behavior in the SAA paradigm. Our results also add to a growing body of evidence on the neural substrates of SAA (Boeke, Moscarello, LeDoux, Phelps, & Hartley, 2017; Bravo-Rivera, Roman-Ortiz, Brignoni-Perez, Sotres-Bayon, & Quirk, 2014; Choi, Cain, & LeDoux, 2010; Kamin, Brimer, & Black, 1963; Lázaro-Muñoz, LeDoux, & Cain, 2010; Martinez et al., 2013; Mineka, 1979; Moscarello & LeDoux, 2013) by demonstrating a dissociation between DH and VH in the contextual regulation of the avoidance response. Intriguingly, the primary function of VH in two-way SAA seems to be suppressive, attenuating avoidant behavior outside the training environment. Future work will explore the broader circuit connections of VH with regions such as the amygdala, nucleus accumbens, and prefrontal cortex (Britt et al., 2012; Canteras & Swanson, 1992; Pitkänen, Pikkarainen, Nurminen, & Ylinen, 2000; Risold & Swanson, 1996; van Groen & Wyss, 1990).

